# Tau monomer encodes strains

**DOI:** 10.1101/325043

**Authors:** Apurwa M. Sharma, Talitha L. Thomas, DaNae R. Woodard, Omar M. Kashmer, Marc I. Diamond

**Affiliations:** Center for Alzheimer’s and Neurodegenerative Diseases, University of Texas Southwestern Medical Center, Dallas, TX, USA; Graduate Program in Biochemistry, Division of Biology and Biomedical Sciences, Washington University in St. Louis, St. Louis, MO, USA

## Abstract

Tauopathies have diverse presentation, progression, and neuropathology. They are linked to tau prion strains, self-replicating assemblies of unique quaternary conformation. Strains can be propagated indefinitely in cultured cells, and induce unique patterns of transmissible neuropathology upon inoculation into mice. Aggregates from a single strain reproduce only that strain upon re-inoculation into cells or mice. DS9 and DS10 cell lines propagate distinct synthetic strains. Surprisingly, DS9 monomer inoculated into naïve cells encoded an identical “sub-strain,” whereas DS10 monomer encoded multiple sub-strains. Sub-strains produced distinct pathology upon inoculation into a tauopathy mouse model (PS19). Brain-derived tau monomer from an Alzheimer’s brain encoded a single strain. Monomer from a corticobasal degeneration brain encoded three sub-strains in which monomer from each encoded all three upon re-inoculation into cells. Tau monomer thus adopts multiple, stable seed-competent conformations, each of which encodes a limited number of strains. This provides insights into the origins of distinct tauopathies.

## Introduction

Tauopathies are a diverse group of neurodegenerative diseases defined by the accumulation of tau amyloids in neurons and glia (1). They include Alzheimer’s disease (AD) and corticobasal degeneration (CBD), among many clinical and neuropathological syndromes. Most are sporadic, and some are caused by dominant mutations in the microtubule associated protein tau (MAPT) gene (1). The cause of diversity in sporadic tauopathies is poorly understood. We initially hypothesized that the diversity of human tauopathies might arise from distinct, self-propagating assemblies based on our observation that a single tau monomer would stably propagate distinct structures *in vitro*, depending on the template to which it was exposed (2). Experiments from our group and others subsequently indicated that tau transmits amyloid pathology into cultured cells and mouse brain that can move from cell to cell (3,4), and led to the idea that it was “prion-like.” Prions form “strains,” which are distinct amyloid structures derived from a single protein that propagate indefinitely in living systems, and which underlie diverse and predictable patterns of neuropathology in humans and mice.

We found that tau forms strains that can be propagated from cell-to-cell in a stable cell line that express the repeat domain (RD) containing two disease-associated mutations (P301L/V337M) fused to yellow fluorescent protein (RD-YFP). We initially created two “artificial” strains based on inoculation of recombinant fibrils into RD-YFP cells and isolation of two clonal lines that stably propagated aggregates of distinct conformation: DS9 and DS10. After inoculation into a mouse model of tauopathy (PS19) that is based on expression of 1N4R human tau with a single disease-associated mutation (P301S) (5), we observed that DS9 and DS10 strains created distinct neuropathological patterns. These could be passaged stably across multiple generations of mice, and finally back into the RD-YFP biosensor cells, where they displayed their original phenotype(6). Additionally, in this work we isolated distinct tau strain ensembles from the brains of patients with different tauopathies, which linked different strains to various neuropathological syndromes. Based on these data we proposed that tau should be considered a prion (6).

Subsequently, we created 18 distinct tau prion strains and observed that following intracerebral inoculation they created distinct pathological patterns and rates of progression. Indeed, others had previously made similar observations of neuropathological diversity upon inoculation of tau fibril preparations from human disorders(7). We have concluded that, in addition to probable contributions of environmental and genetic factors, the diversity of tauopathy might be explained in large part by the strains that underlie them, in other words, by protein conformation.

Our recent work indicates that tau monomer adopts multiple, stable structures that we have grouped into two general categories: “inert” (M_i_) or “seed-competent” (M_s_) (Mirbaha et al., eLife, 2018, *in press*). Whereas M_i_ does not spontaneously self-assemble or seed aggregation, M_s_ has a distinct conformation that allows self-assembly and seeding activity (Mirbaha et al., eLife, 2018, *in press*). Structural studies suggest that this is based on exposure of critical amino acids that have previously been determined to mediate amyloid formation (VQIINK/VQIVYK) {Von Bergen et.al. 2000}{Von Bergen et.al‥ 2001}. In M_s_ these amino acids are predicted to be solvent-exposed, while in M_i_ they predicted to be buried in hairpins and rendered relatively inaccessible for inter-molecular interactions. We have previously isolated M_s_ from recombinant sources and human brain (Mirbaha et al., eLife, 2018, *in press*).

The existence of M_s_ as a unique conformational structure raised the question of what is the role of monomer in strain formation. We have imagined two possibilities. First, a single structure of M_s_ might underlie multiple distinct assemblies. This has been suggested as the basis of two different fibril morphologies isolated from an AD patient, in which the same essential monomeric building block appears to constitute the “core” of the amyloid of both structures (8). Second, an M_s_ protein might adopt multiple distinct conformations as an ensemble, each producing a single strain or subset of strains upon self-association as a multimeric assembly. This would predict that the diversity of tau prion strains we have previously described (6,9) could be linked back to a distinct set of conformers of tau monomer. We have addressed this question by studying strains propagated in HEK293T cells, DS9 and DS10, that were derived from recombinant fibrils, and others derived from AD and CBD brain.

## RESULTS

### Tau monomer encodes restricted strain information

We previously characterized in detail two tau strains, DS9 and DS10{Sanders:2014cv}. We hypothesized that, if M_s_ acts as an “unrestricted” building block, monomer derived from either DS9 or DS10 would produce a diversity of strains. We isolated total lysate or monomer from DS9 using size exclusion chromatography (SEC) or by passage through a 100kD size-exclusion filter, methods that we have previously determined to faithfully exclude larger assemblies (10)(Mirbaha et al., eLife, 2018, *in press*). We then inoculated DS1 cells, which lack any aggregates, and used FACS to isolate single aggregate-containing cells, with which we established monoclonal lines for further analysis. We initially used confocal microscopy to characterize the various colonies for inclusion morphology, which serves as a rough surrogate for strain identity(6) (9). For M_s_ derived from DS9, we observed no variance—each of 52 clones exhibited the speckled conformation previously observed for DS9(6) (Fig. 1A). We isolated 6 typical “sub-strains,” termed DS9.1-9.6, for further characterization in detail. In addition, we passaged the lysate through a 100kDa size exclusion filter and used it to inoculate DS1 cells, creating 31 monoclonal lines. All 31 lines were indistinguishable from DS9 by confocal microscopy. Application of unfractionated DS9 lysate, or lysate of any of the sub-strains produced a single population of clones, all identical to DS9 in morphology (Fig. 1A).

**Figure 1.**
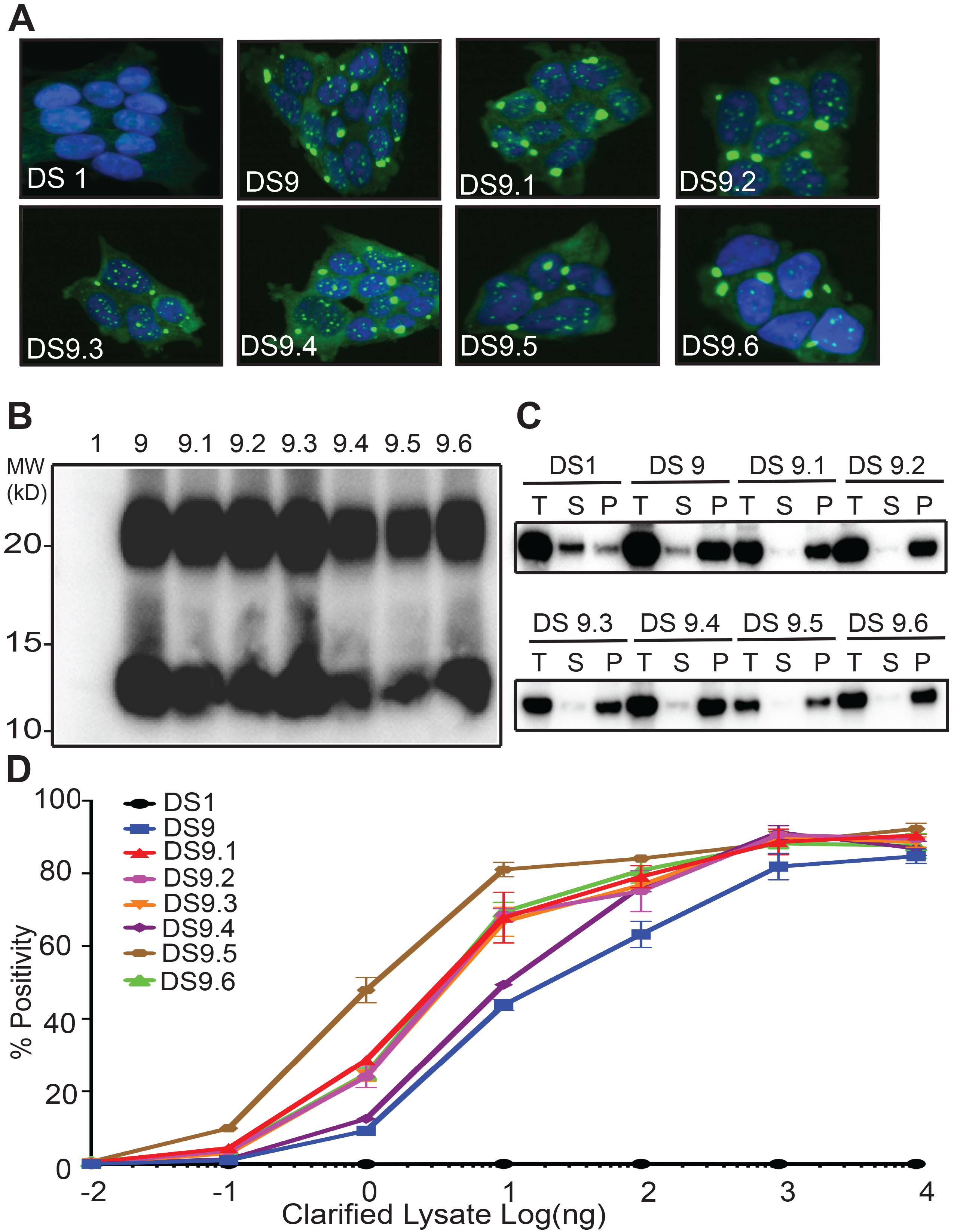
Tau M_s_ from DS9 retains strain identity. (A) Clones isolated from DS9 monomer (9.1-9.6) show similar morphological characteristics to DS9. (B) Limited proteolysis digests all the monomer from Clone 1, but reveals similar protease resistant band patterns for DS9 and DS9.1-9.6. Both DS9 and its sub-strains exhibited a major band around 10 kDa, and a larger band between 20 and 25 kDa. (C) Sedimentation analysis was performed on DS1, 9, and its substrains DS9.1-9.6. Pellet (P) was isolated from supernatant (S) by ultracentrifugation. For all the samples, supernatant to pellet ratio on the gel was 1:1. DS1 had tau in the supernatant, whereas DS9 and its substrains had tau predominantly in the pellet. (D) DS9 and sub-strains had similar seeding activities upon transduction into P301S FRET biosensor cells.

We have previously used proteolysis of insoluble tau to discriminate distinct strains. This reveals variation in the protease resistant “cores” of tau aggregates (6) (9). DS1 had no protease resistant species. DS9 and DS9.1-9.6 exhibited very similar limited proteolysis patterns, with a band at 10 kDa, and a second strong band between 20 and 25 kDa (Fig. 1B). Next we used sedimentation analysis to differentiate the clones by subjecting the clarified lysate to high-speed centrifugation to separate the soluble from insoluble species. DS9 and DS9.1-9.6 exhibited similar fractionation patterns, with most tau being insoluble (Fig. 1C). Finally, we used an established biosensor cell line that expresses tau RD (P301S) fused to cyan and yellow fluorescent proteins (RD-CFP/YFP) to monitor the ability of strains to trigger intracellular aggregation. DS9 and DS9.1-9.6 had identical maximal seeding, with a slight variance in the dose responses (all within ~1 log of concentration, which is typical for independent isolates; Fig. 1D). Thus, monomer from DS9 faithfully encoded 6 identical DS9 sub-strains.

### DS10 monomer produces multiple sub-strains

Extending our studies with DS9, we isolated total lysate or M_s_ from DS10, transduced DS1 cells, and isolated multiple monoclonal inclusion-bearing lines. As we have previously reported, application of unfractionated DS10 lysate produced a single population of clones, all identical to DS10 (data not shown) (6). However DS10 monomer created five distinct sub-strains, easily discerned by inclusion morphology: 36% were ordered (and indistinguishable to the parent strain) termed DS10.1; 21% were speckled, termed DS10.2; 13% thread-like, termed DS10.3; 9% disorganized, termed DS10.4 (Fig. 2A, Table 1). Approximately 21% of cells formed a fifth strain, DS10.5, that rapidly “sectored,” i.e. lost its inclusions, and thus could not be further characterized. All strain images were analyzed by a blinded reviewer (D.R.W.). DS10 monomer was also isolated using a 100kDa cut-off filter. After transduction into DS1 cells we isolated monoclonal lines, and obtained similar strain diversity (Table 2).

**Figure 2:**
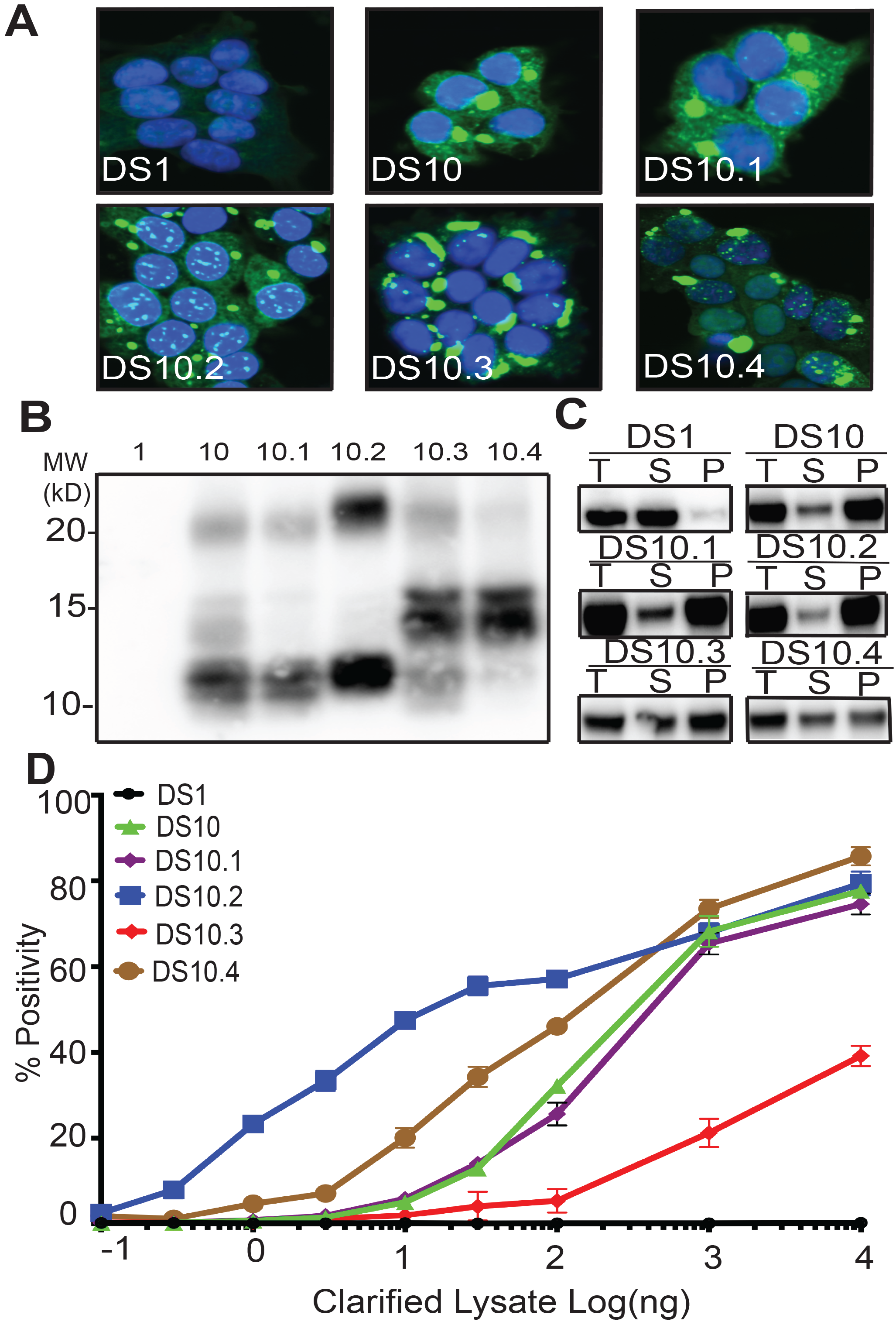
Tau M_s_ from DS10 creates multiple sub-strains. (A) Clones isolated from DS10 monomer give rise to cells with multiple morphologies. Four sub-strains were discriminated based on multiple tests. (B) Limited proteolysis of RD-YFP using pronase differentiated the protease resistant cores in the sub-strains. (C) Sedimentation analysis of RD-YFP was performed on DS1, 10, and DS10.1-4. Pellet (P) was isolated from supernatant (S) by ultracentrifugation. For all the samples, supernatant to pellet ratio on the gel was 1:1. DS1 had RD-YFP in the supernatant; DS10, 10.1 and 10.2 had most RD-YFP in the pellet; DS10.3 and DS10.4 had mixed RD-YFP solubility. (D) DS10 sub-strains had distinct seeding activities upon transduction into P301S FRET biosensor cells.

**Table 1:**
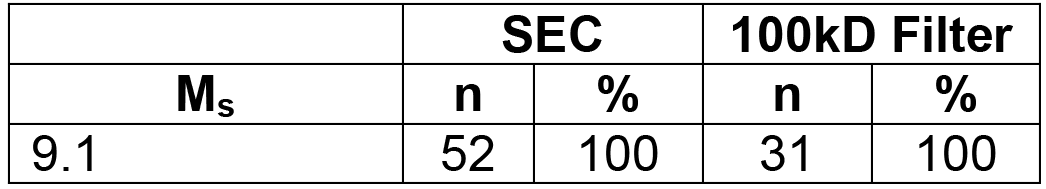
Sub-strains generated from DS9 monomer isolated by SEC or cutoff filter. Tau RD-YFP monomer (M_s_) was isolated from DS9 by either by SEC or 100kD cutoff filter and inoculated into DS1 to create sub-strains. Multiple clones were isolated and characterized by morphology. Columns indicate the number of clones identified (n) and the percentage this represents of the total (%). A single sub-strain was observed regardless of purification method.

| **M<sub>s</sub>** | **SEC** |  | **100kD Filter** |  |
| --- | --- | --- | --- | --- |
|  | **n** | **%** | **n** | **%** |
| 9.1 | 52 | 100 | 31 | 100 |

**Table 2:** Sub-strains generated from DS10 monomer isolated by SEC or cutoff filter. Tau RD-YFP monomer (M_s_) was isolated from DS10 by either by SEC or 100kD cutoff filter and inoculated into DS1 to create sub-strains. Multiple clones were isolated and characterized by morphology. Columns indicate the number of clones identified (n) and the percentage this represents of the total (%). Isolation of M_s_ from DS10 by SEC or 100kD each enabled a similar proportion of sub-strains to form.

| **M<sub>s</sub>** | **SEC** |  | **100kD Filter** |  |
| --- | --- | --- | --- | --- |
|  | **n** | **%** | **n** | **%** |
| 10.1 | 19 | 36 | 12 | 43 |
| 10.2 | 11 | 21 | 3 | 11 |
| 10.3 | 7 | 13 | 5 | 20 |
| 10.4 | 5 | 9 | 3 | 11 |
| 10.5 (sectored) | 11 | 21 | 4 | 15 |
| **Total** | **53** | **100** | **27** | **100** |

We used limited proteolysis to compare the four monoclonal DS10 sub-strains (Fig. 2B). DS10 and DS10.1 exhibited similar protease resistant doublets around 10 kDa and a relatively light band around 20 kDa. DS10.2 exhibited a single strong band at 10 kDa and a strong band between 20 and 25 kDa. DS10.3 and 10.4 exhibited a distinctive strong band at 15 kDa. DS10.3 exhibited a band at 10 kDa, which was mostly absent in DS10.4. Sedimentation analysis helped further discriminate the sub-strains: DS10, DS10.1 and DS10.2 primarily manifested an insoluble fraction and a small soluble fraction. By contrast, DS10.3 and DS10.4 exhibited mixed solubility, with about half soluble and insoluble tau (Fig. 2C). DS10 sub-strains also had very different seeding efficiency that spanned >2 log orders of concentration (Fig. 2D). DS10 and DS10.1 had almost identical seeding profiles, with EC_50_ of 300 ng of clarified lysate. DS10.2 had an EC_50_ of 10 ng. DS10.3 had a much weaker seeding activity with EC_50_ at >10 μg (unable to be determined accurately), and reduced maximal seeding efficiency at ~50%. DS10.4 had an EC_50_ of 100 ng. Based on multiple measures, we concluded that unlike DS9, DS10 monomer encoded distinct sub-strains, each with unique morphological and biochemical characteristics.

### Conformational transitions of tau monomer

Sub-strains of DS10 could arise because multiple distinct monomers co-exist, each constrained to form only a single sub-strain. In this case, M_s_ from each sub-strain would be predicted to recreate that same sub-strain (as for DS9.1-9.6). Alternatively, M_s_ derived from DS10 might exist as a conformationally restricted ensemble, with relatively low kinetic and energetic barriers between various states that become “locked in” upon assembly as a fibril. This led us to predict that monomer from any sub-strain could establish all of the others upon re-inoculation. To test these ideas, we isolated monomer from the various sub-strains, re-introduced them into DS1, and isolated n=47-51 monoclonal cell lines bearing inclusions. We analyzed each of these using blinded scoring of intracellular inclusion morphology. In 49 sub-strains derived from DS10.1 monomer, 45 appeared identical to DS10.1, except for 4 that formed DS10.5 (sectored) (Table 3). This suggested M_s_ from DS10.1 was relatively restricted in its conformation. We obtained very different results with the other DS10 sub-strains. M_s_ derived from each of DS10.2, DS10.3, or DS10.4 gave rise to the other three sub-strains including DS10.1, with predomination of the original sub-strain (Table 3). This indicated that M_s_ from these DS10 substrains existed as conformational ensemble. Taken together our experiments indicated that while multimeric assemblies from total lysate exhibit consistent strain behavior in all cases, i.e. faithful replication, tau monomer (M_s_) is less reliable. While M_s_ derived from DS9 was completely restricted to a single strain, M_s_ from DS10.1 predominantly encoded one sub-strain, and others (DS10.2, DS10.3, DS10.4) adopted a defined set of strains that could give rise to all four. Every DS10 substrains gave rise to the sectored population.

**Table 3:**
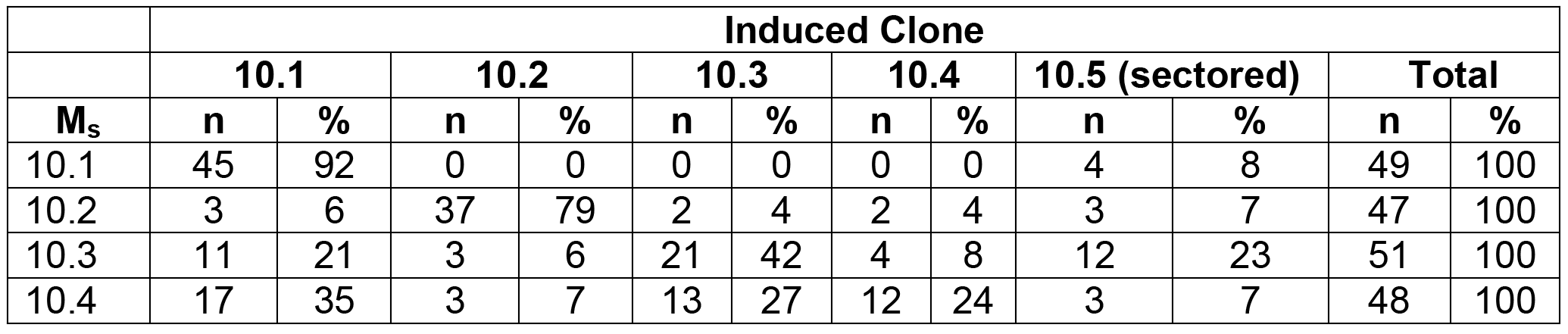
Quantification of second generation of sub-strains obtained from DS10. M_s_ from each sub-strain of DS10 (10.1-10.5) was inoculated into DS1, and clones of the induced strains were characterized. DS10.1 largely produced a single predominant strain identical to DS10.1 (92%) and another strain DS10.5 that rapidly sectored (8%). DS10.2-10.4 each recreated the other strains, with the exception of DS10.1. Columns indicate the number (n) of clones characterized and the percentage of the total (%) in each case.

| **M<sub>s</sub>** | **Induced Clone** |  |  |  |  |  |  |  |  |  |  |  |
| --- | --- | --- | --- | --- | --- | --- | --- | --- | --- | --- | --- | --- |
|  | **10.1** |  | **10.2** |  | **10.3** |  | **10.4** |  | **10.5 (sectored)** |  | **Total** |  |
|  | **n** | **%** | **n** | **%** | **n** | **%** | **n** | **%** | **n** | **%** | **n** | **%** |
| 10.1 | 45 | 92 | 0 | 0 | 0 | 0 | 0 | 0 | 4 | 8 | 49 | 100 |
| 10.2 | 3 | 6 | 37 | 79 | 2 | 4 | 2 | 4 | 3 | 7 | 47 | 100 |
| 10.3 | 11 | 21 | 3 | 6 | 21 | 42 | 4 | 8 | 12 | 23 | 51 | 100 |
| 10.4 | 17 | 35 | 3 | 7 | 13 | 27 | 12 | 24 | 3 | 7 | 48 | 100 |

### Sub-strains induce distinct pathologies in PS19 mice

*Bona fide* tau prion strains create distinct and predictable pathologies *in vivo*. We previously used inoculation into PS19 mice that express full-length (1N,4R) human tau with a P301S mutation(5) to study the pathology of strains (6,9). To test the activities of sub-strains, we inoculated n=5 replicates of DS9, DS9.1, DS10, DS10.1-10.4 into the left hippocampus of PS19 mice at 3mos. Mice were age-and gender-matched to the extent possible, and derived from multiple independent litters. 4 weeks after inoculation we analyzed the brains of the mice using AT8 antibody, which recognizes p-Ser202 and p-Thr205 of tau (Fig. 3).

**Figure 3:**
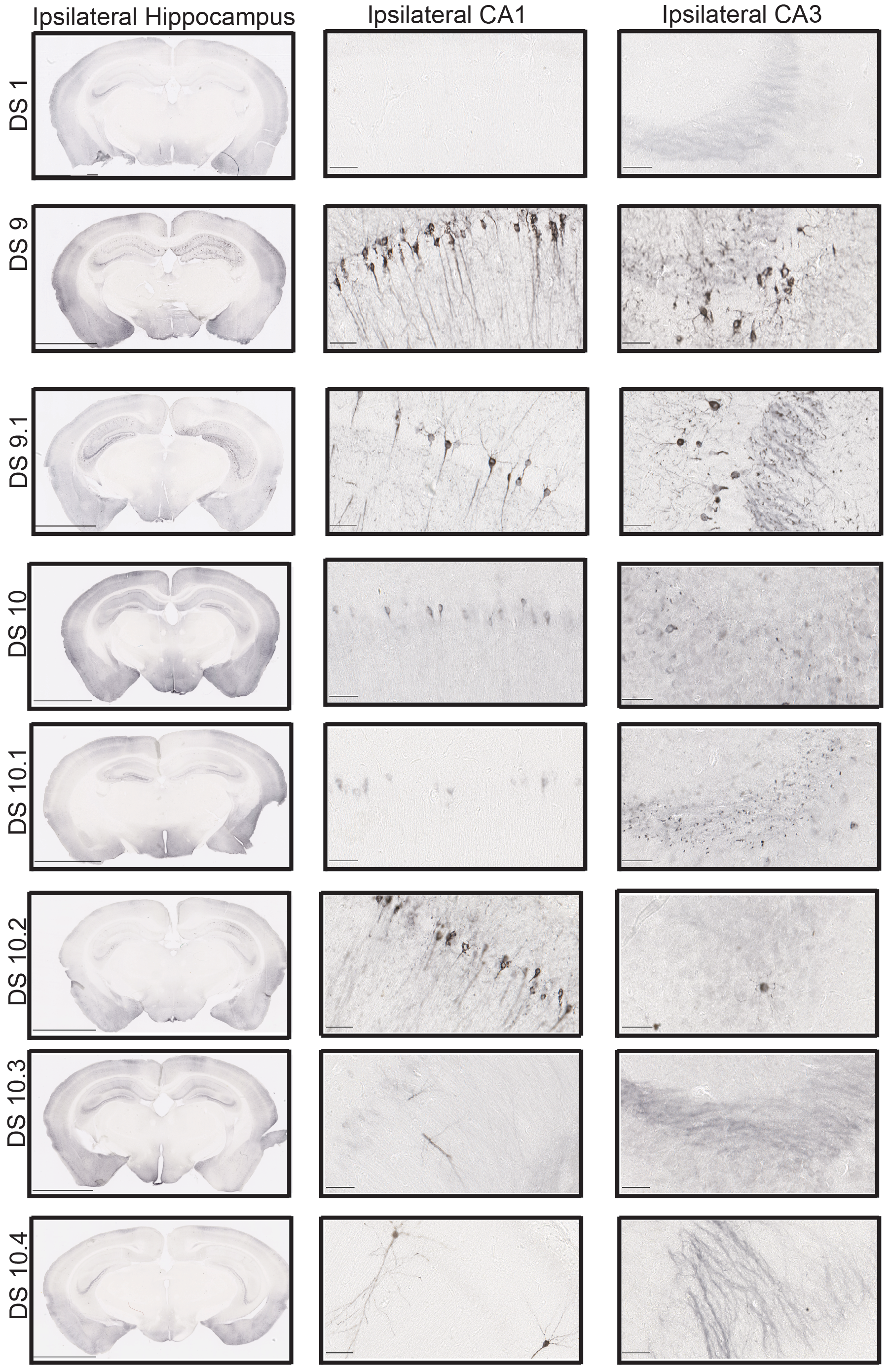
Sub-strains trigger unique tau pathology in P301S mice. 10ug of clarified lysate was injected into the hippocampi of 3 mo P301S mice, followed by AT8 staining after 4 weeks. DS9 and DS9.1 induced similar pathology in the CA1 and CA3 regions. DS1 induced no pathology. DS10 and DS10.1 induced similar pathology in CA1 and CA3. In both cases, we observed AT8 staining in the cell body throughout CA1 and AT8-positive puncta throughout CA3. DS10.2 induced AT8 signal in both the cell body and along the axons in CA1. There was very little pathology in CA3. DS10.3 induced very little AT8 signal in CA1 and none in CA3. DS10.4 likewise induced little pathology in CA1 region and none in CA3.

Mice inoculated with DS1 controls exhibited no tau pathology in the hippocampus, CA1, or CA3. DS9, DS10, and all sub-strains induced different types of pathology. DS9 and DS9.1 induced tangle-like inclusions throughout CA1 and CA3 in patterns that were indistinguishable from one another, consistent with prior observations (6). Also as expected, DS10 induced AT8-positive puncta in mossy fiber tracts of the hippocampus with strong staining patterns in the cell body, and AT8 positive puncta in CA3. DS10.1 induced pathology similar to DS10 in both CA1 and CA3. DS10.2 induced tangle-like inclusions throughout CA1, but with little or no pathology in CA3. DS10.3 had very low seeding efficiency and induced pathology only in CA1. Clone 10.4 also had low seeding and induced pathology only in CA1. A blinded reviewer (O.M.K.) analyzed slides derived from all inoculations (n=50), and classified them by group. DS9 and DS9.1 were indistinguishable. DS10.1-10.3 were all easily distinguished, with a single error for DS10.4, which was misclassified in one instance (Table 4).

**Table 4:** Analysis of AT8 signal in hippocampi of injected mice. PS19 mice were inoculated with sub-strains from DS9 and DS10 at 3 mo into the hippocampus (n=5 each). After 8 weeks, coronal sections of hippocampus were analyzed by a blinded reviewer educated on different sections as to the characteristics of each clone. One error occurred in analysis of 40 brains.

| Strain Inoculated | Blinded Analysis |
| --- | --- |
| DS9 | DS9 and DS9.1 were indistinguishable. |
| DS9.1 |  |
| DS10 | DS10 and DS10.1 were indistinguishable. |
| DS10.1 |  |
| DS10.2 | DS10.2 were correctly classified |
| DS10.3 | DS10.3 were correctly classified. |
| DS10.4 | DS10.4 correctly classified 4/5 times, 1/5 incorrectly classified as DS10.3 |

### Distinct M_s_ conformations in AD and CBD brains

The preceding studies with DS9 and DS10 were based on tau RD-YFP fusions, and left uncertain whether tau monomer derived from human tauopathies would similarly encode strains. We have previously determined strain composition in AD and CBD patient brains to be quite distinct (6). Thus we used these syndromes as a source of tau monomer. We gently lysed the brains by dounce homogenization, a method previously determined not to liberate M_s_ from pre-existing fibrils (Mirbaha, eLife 2018, *in press*), and used an anti-tau antibody directed at the N-terminus (HJ8.5) to purify full-length tau and resolve monomer by SEC. We inoculated DS1 cells with total lysate from an AD patient and recovered a single clonal morphology, AD(t). We also used AD monomer isolated by SEC to inoculate DS1 cells and recovered an identical clonal morphology, AD(m). As we had previously observed(6), each clone exhibited a single “speckled” conformation, indicating that we had most likely propagated a single strain (Fig. 4A). We also isolated AD monomer by passage through a 100k kDa cut-off filter and inoculated DS1 cells, isolating 23 monoclonal lines. All clones had similar morphological characteristics to AD(t) and AD(m) (Table 5). To better characterize the tau prions propagated in these clones, we performed limited proteolysis on the lysates from AD(t) and AD(m) clones. This revealed identical digestion patterns (Fig. 4B). We also compared them using sedimentation analysis. Strains derived from AD(m) or AD(t) consisted of both soluble and insoluble protein (Fig. 4C). Seeding analyses from the cell lines also revealed very similar potencies (Fig. 4D). We concluded that tau M_s_ from the AD patient encoded the same single strain as total lysate.

**Table 5:** Sub-strains generated from AD monomer isolated by SEC or cutoff filter. Tau monomer (M _s_) from AD brain was purified by immunoprecipitation followed by SEC or passage through a 100kD cutoff filter, prior to inoculation into DS1 cells. Columns indicate the number of clones identified (n) and the percentage this represents of the total (%). A single AD sub-strain was identified regardless of purification method.

|  | SEC |  | 100kD Filter |  |
| --- | --- | --- | --- | --- |
| M<sub>s</sub> | n | % | n | % |
| AD(m) | 47 | 100 | 23 | 100 |

**Figure 4:**
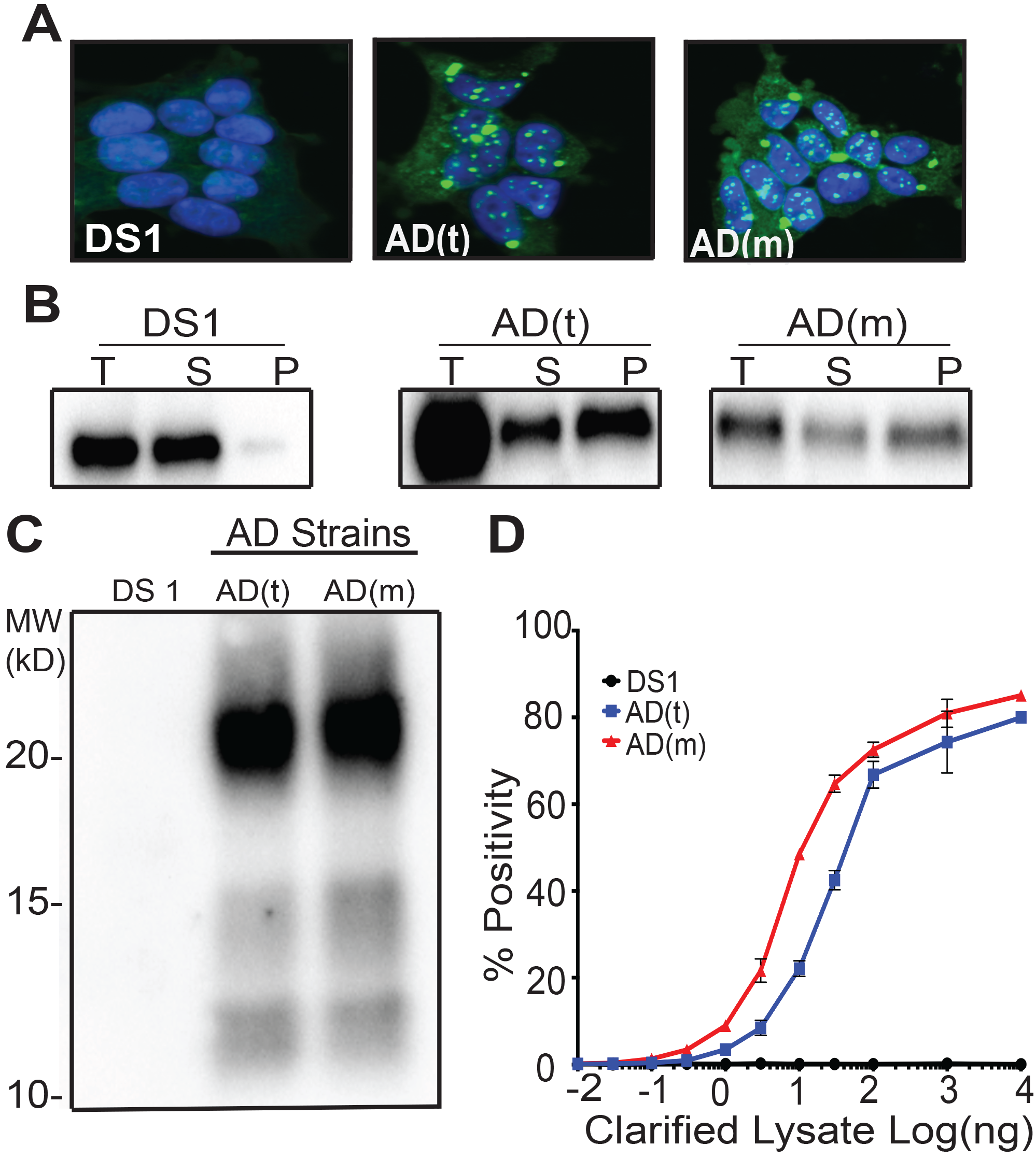
M_s_ derived from an AD patient produces a single strain. (A) Clonal cell lines derived from AD-derived total lysate, AD(t), and M_s_, AD(m) exhibited identical inclusion morphologies. (B) RD-YFP derived from AD(t) and AD(m) had similar solubility profiles. RD-YFP in DS1 was present in the supernatant fraction only, as expected. (C) Limited proteolysis of RD-YFP aggregates from AD(t) and AD(m) clonal lines produced identical band patterns. (D) Lysates of AD(t) and AD(m) clonal lines had similar seeding profiles.

Next we evaluated strains derived from total lysate or monomer from a CBD patient. Upon transduction of the DS1 cells, the total CBD lysate produced two distinct strains: CBD1(t) and CBD2(t). M_s_ from the CBD brain created two strains identical to those derived from total lysate CBD1(m) and CBD2(m), plus a third strain, CBD3(m) (Fig. 5A, Table 6). CBD monomer was also isolated using 100k kDa cut-off filter and transduced into DS1 line to produce distinct clones. Based on morphology, the proportion of sub-strains was similar to that obtained using gel filtration (Table 6). Whether derived from total lysate or monomer, CBD1(t) and CBD1(m), or CBD2(t) and CBD2(m) exhibited similar sedimentation properties (Fig. 5B), proteolysis patterns (Fig, 5C), and seeding activities (Fig 5D). CBD3(m) exhibited a unique morphology, with ordered aggregates (Fig. 5A), distinct sedimentation (Fig. 5B), proteolysis (Fig. 5C), and higher seeding activity (Fig 5D).

**Table 6:** Sub-strains generated from CBD monomer isolated by SEC or cutoff filter. M_s_ from CBD brain was purified by immunoprecipitation followed by SEC or passage through a 100kD cutoff filter, prior to inoculation into DS1 cells. CBD sub-strains, CBD1-3(m), were quantified. Isolation of M_s_ from CBD brain by SEC or cutoff filter enabled a similar proportion of sub-strains to form. Columns indicate the number of clones identified (n) and the percentage this represents of the total (%). M_s_ created similar strain patterns regardless of filtration method.

|  | SEC |  | 100kD Filter |  |
| --- | --- | --- | --- | --- |
| M<sub>s</sub> | n | % | n | % |
| CBD1(m) | 20 | 36 | 4 | 22 |
| CBD2(m) | 18 | 33 | 7 | 39 |
| CBD3(m) | 17 | 31 | 7 | 39 |
| **Total** | **55** | **100** | **18** | **100** |

**Figure 5:**
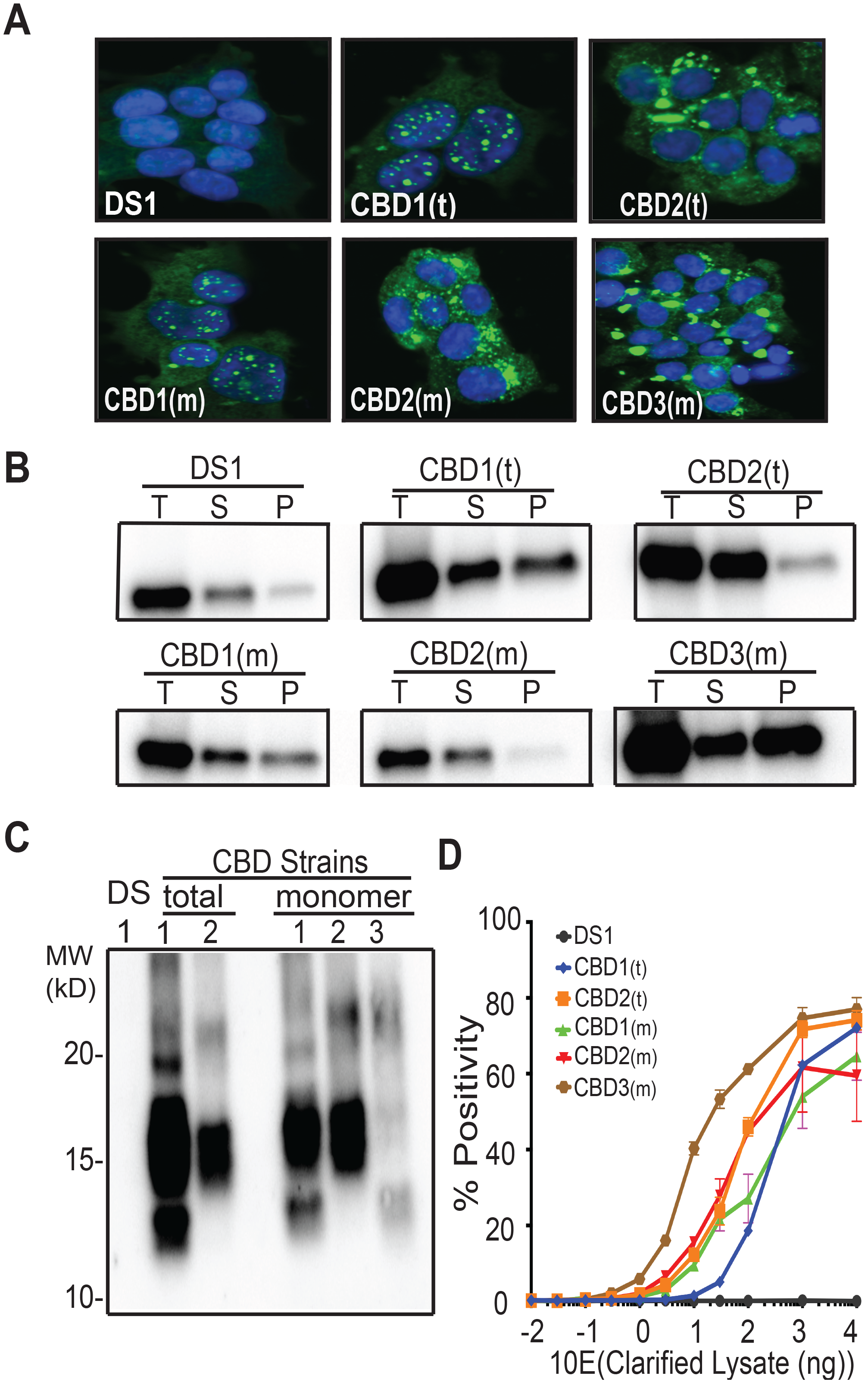
M_s_ derived from a CBD patient produces three distinct strains. (A) Clonal lines derived from CBD total lysate had two distinct inclusion patterns, CBD1(t), CBD2(t). Clonal lines derived from CBD M_s_ had two patterns identical to those from the total lysate, CBD1(m), CBD2(m), and a third, CBD3(m). (B) RD-YFP from CBD1(t) and CBD1(m) had mixed solubility. In CBD2(t) and CBD2(m), most RD-YFP was present in the supernatant. CBD3(m) had mixed solubility with most RD-YFP in the insoluble fraction. (C) RD-YFP aggregates from CBD1(t) and CBD1(m) exhibited similar patterns of proteolysis, with protease resistant bands around 10-15 kDa and a strong band around 15 kDa. Proteolysis of RD-YFP from CBD2(t) and CBD2(m) only produced a strong band around 15 kDa. CBD3(m) had a unique proteolysis pattern with a band around 10-15 kDa. (D) Lysates from clones CBD1(t), CBD1(m), CBD2(t) and CBD2(m) had similar seeding profiles, while lysate from CBD3(m) had more potent seeding activity.

We concluded that M_s_ from CBD could be comprised of three independent seed-competent monomers, or that a single monomer exhibited conformational “flexibility” before forming larger assemblies of a particular structure. In the first case, we predicted that monomer from each of the three CBD sub-strains would only produce the parent strain. Conversely, if a single monomer could form three different strains, then monomer from any one of the CBD sub-strains would recreate the panel. We isolated monomer from the three CBD sub-strains, re-inoculated DS1, and evaluated the resultant clones by morphology. M_s_ derived from each sub-strain CBD1,2,3 preferentially encoded the parent strain, but also the other two (Table 7). These data indicated that M_s_ derived from CBD represented a conformational ensemble (analogous to DS10) that encoded a defined subset of strains. Conversely, M_s_ derived from the AD patient had a more restricted conformational state that only encoded a single strain.

**Table 7:** Quantification of strains derived from CBD-derived sub-strains. Monomeric RD-YFP derived from each CBD sub-strain was used to inoculate DS1. The resultant clones were then characterized by morphology. M_s_ from each sub-strain recreated all three, with a preference for the strain of origin. Columns indicate the number of clones identified (n) and the percentage this represents of the total (%).

|  | Induced Clone |  |  |  |  |  |  |  |
| --- | --- | --- | --- | --- | --- | --- | --- | --- |
|  | CBD1 |  | CBD2 |  | CBD3 |  | Total |  |
| Input M<sub>s</sub> | n | % | n | % | n | % | n | % |
| CBD1(m) | 30 | 57 | 7 | 15 | 10 | 20 | 47 | 100 |
| CBD2(m) | 9 | 17 | 29 | 62 | 10 | 20 | 48 | 100 |
| CBD3(m) | 13 | 26 | 11 | 23 | 31 | 60 | 55 | 100 |

### Discussion

We have previously defined two conformational ensembles of tau: M_i_, which is inert, and M_s_, which is competent for self-assembly and seeding (Mirbaha et al., eLife 2018, *in press*). It has been unclear to what extent M_s_ encodes strain information. We now conclude that M_s_ represents an ensemble of structures that each encodes a restricted set of strains. M_s_derived from artificially-derived strains (DS9 and DS10) replicated the parent strain exclusively (DS9), or a series of sub-strains, one of which closely resembled the parent strain (DS10). These sub-strains each produced unique pathology upon inoculation into the PS19 tauopathy mouse model. Turning to patient-derived M_s_, we observed that, like total brain lysate, monomer from an AD patient encoded a single strain. Conversely, while total lysate from a CBD patient encoded two strains, M_s_ derived from CBD lysate encoded these two strains, plus a third. Thus we hypothesize that tau, despite being unstructured by classical biophysical measures, can adopt multiple, stable conformational ensembles in its seed-competent form, M_s_. These may restrict subsequent assembly to a single fibril conformation (as for M_s_ derived from DS9 or AD), or may enable formation of a limited number of assembly structures that constitute individual strains (as for M_s_ derived from DS10 and CBD).

Intriguingly, monomer from DS10 produced five sub-strains. DS10.1 resembled DS10 in every respect, except that monomer from this sub-strain did not recreate the original diversity seen in monomer from DS10. This suggests that in forming DS10.1, M_s_ adopts a structure very similar to that of DS10 monomer, but with a higher energetic or kinetic barrier that restricts its conformation. It appears that in isolating M_s_ from DS10 we enabled a new, restricted M_s_ conformation to emerge and be trapped within the DS10.1 strain. Nonetheless, the overall similarity of DS10 and DS10.1 in terms of inclusion morphology, seeding activity, proteolytic patterns, and neuropathology suggests that the core amyloid conformations are likely to be very similar.

A major caveat of this work is that we amplified strains using tau RD-YFP containing two mutations (P301L/V337M), not full-length protein. We recognize that our system may bias towards detection of some strains. It remains to be determined (and studies are ongoing) whether RD-YFP captures the critical core structures of fibrils that occur in patients. For example, prior work by Fitzpatrick et.al. indicated that two distinct fibril morphologies (paired helical filaments and straight filaments) could be detected in AD brain(8). These appeared to derive from different configurations of a single monomer template. This is consistent with our prior work, where 2/6 AD patients we analyzed for strain composition showed presence of two distinct strains, whereas in 4/6 patients, we observed a single strain(6). Nonetheless, based on our characterization of strains derived from M_s_ extracted from the AD and CBD brains, we feel confident in our primary conclusions. In this regard, we were excited to see recent work by Ohhashi et.al, who found that the intrinsically disordered region of Sup35 harbored local compact structure, and also that distinct forms of monomer could give rise to unique Sup35 amyloid conformations(11).

Based on this work, we predict that related “families” of strains will be described in patients, linked hierarchically by structure (Fig. 6). More specifically, we hypothesize that strain families will utilize distinct combinations of amino acids to form “core” structures that are largely preserved even within a monomer. Similarly seed competent monomers will thus adopt a series of structurally related conformations, which can further assemble into amyloid fibrils. Once a multimeric assembly forms, our observations suggest that it replicates relatively faithfully as a strain. This model explains how a single patient might harbor multiple strains, each potentially derived from a topologically restricted ensemble of tau monomer, M_s_. By extension, each distinct tauopathy may develop based on emergence of a restricted conformation of M_s_. This could arise from stochastic or environmental effects, or based on post-translational modifications or other interactions that facilitate conversion of M_i_ to a particular ensemble of M_s_ conformations.

**Figure 6:**
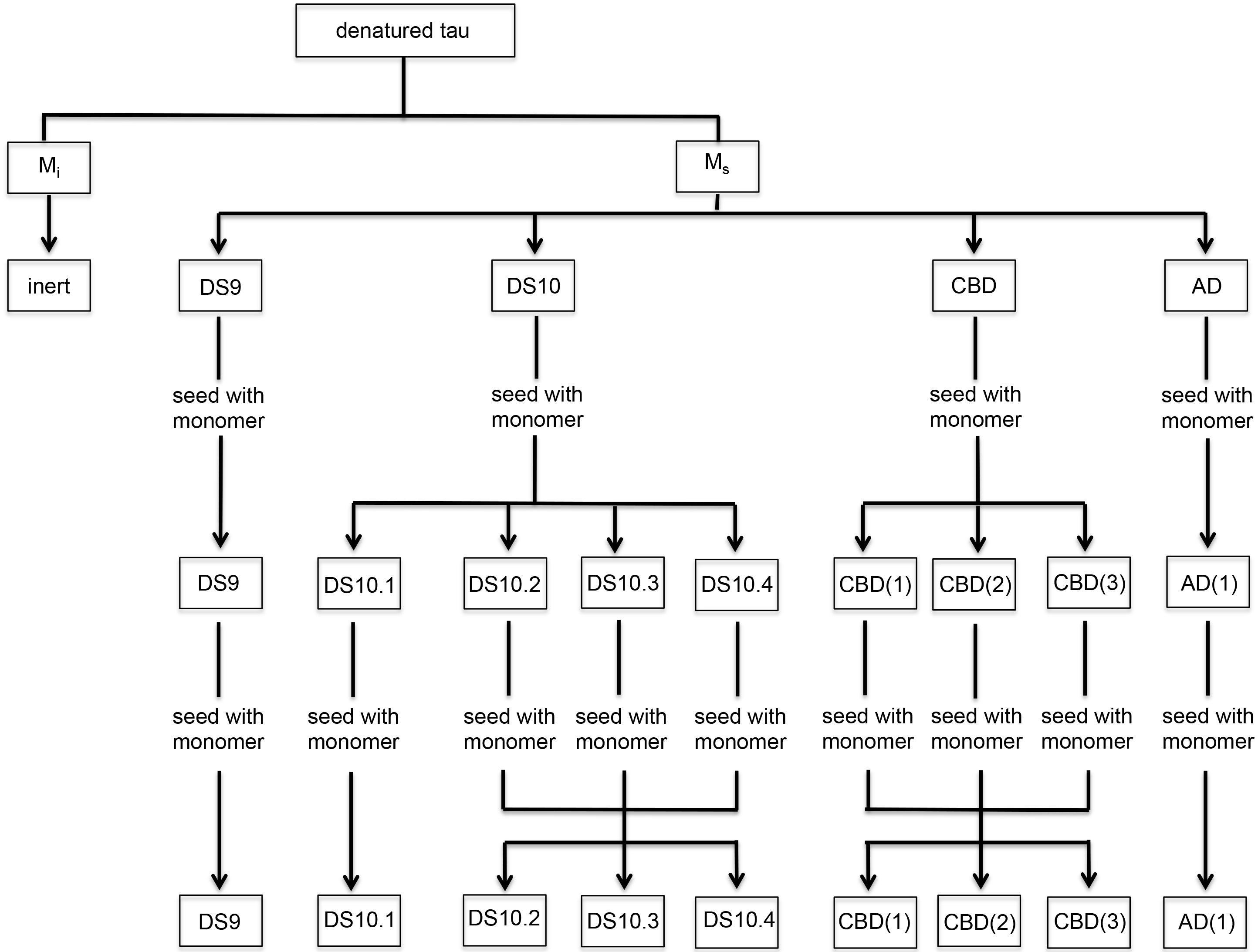
Model for hierarchy of monomer conformations. We propose a hierarchical model that discriminates two general conformational ensembles: M_i_ and M_s_. M_i_ is inert, whereas M_s_ has seeding activity. Within M_s_, multiple conformations exist that can encode individual or multiple strains. Once an assembly forms, a strain will be faithfully replicated, however if M_s_ is isolated from the strain, it can assemble to form a defined set of sub-strains.

In summary, we present a model to explain how diverse aggregate structures form and give rise to phenotypic diversity in the tauopathies. This predicts a hierarchy of restricted M_s_ structural ensembles with varying degrees of relatedness. Tau prion strains, and the diseases they cause, thus may depend ultimately on the original topological restrictions of M_s_ that give rise to them.

## Materials and Methods

### Cell culture

All cells were grown in Dulbecco’s modified Eagle’s medium (Gibco) supplemented with 10% fetal bovine serum (HyClone), 1% penicillin/streptomycin (Gibco), and 1% Glutamax (Gibco). Cells were maintained at 37°C, 5% CO_2_, in a humidified incubator.

### Liposome-mediated transduction of tau seeds

Stable cell lines were plated at a density of 30,000 cells per well in a 96-well plate. After 18h, at 60% confluency, cells were transduced with protein seeds. Transduction complexes were made by combining [11.75 μL Opti-MEM (Gibco) +0.75 μL Lipofectamine 2000 (Invitrogen) with cell lysate at a total volume of 25μL per well. Liposome complexes were incubated at room temperature for 20min before adding to cells. Cells were incubated with transduction complexes for 48h.

### Monoclonal cell isolation

DS1 cells were treated with clarified lysate and monomer using liposome-mediated transduction of tau seeds. After 48h the cells were harvested and re-suspended in flow buffer (1XHBSS, 1%FBS, 1mM EDTA). Aggregate-containing cells were identified based on their particularly bright YFP signal using FACS Aria II SORP cell sorter. Cells were sorted individually into a 96-well plate and grown until confluency to derive clonal lines. Lines stably maintaining aggregates were moved to larger plates and amplified for further studies.

### Western blot

Cell pellets were thawed on ice, lysed by triturating in PBS containing 0.05% Triton-X and a cOmplete mini protease inhibitor tablet (Roche), and clarified by 5-minute sequential centrifugations at 500 x g and 1000 x g. Total protein concentration of the clarified lysate was measured using Bradford Assay (Bio-Rad). clarified lysate was mixed with 2X SDS buffer (final SDS concentration 4%) and run on NuPAGE 4-12% Bis-Tris Gel at 150V for ~75 min. The gel was then transferred onto Immobilon P membrane for 1hr at 20V using a semi-dry transfer apparatus (Bio-Rad). The membrane was then blocked with 5% non-fat dry milk in TBST for 1h before primary rabbit anti-tau monoclonal antibody (Tau A, which was raised against QTAP…KIGSTENL) was added at 1:1000 and placed in a shaker overnight at 4°C. The membrane was then washed 4 times with TBST at 10 min intervals. The membrane was then re-probed with goat anti-rabbit secondary antibody for 1.5h at room temperature. The membrane was then washed 4 times with TBST. Finally the membrane was exposed to ECL Prime western blot detection kit (GE Lifesciences) for 2 min and imaged with a Syngene digital imager.

### Sonication and size exclusion chromatography

Clarified lysate was sonicated using a Q700 Sonicator (QSonica) at a power of 100-110 watt (Amplitude 50) at 4°C for 1h. Samples were then centrifuged at 21000 x g for 10 min and 1 mL of supernatant was loaded into a Superdex 200 Increase 10/300 GL column (GE Healthcare) and eluted in PBS buffer at 4°C. After measuring the protein content of each fraction with a Bradford assay using a plate reader (Tecan M1000), we aliquoted and stored samples at −80°C until further use. Each aliquot was thawed immediately before use. The molecular weight/radius of gyration of proteins in each fraction was estimated by running gel filtration standards (Bio-Rad): Thyroglobulin (bovine) 670 kDa/845nm; γ-globulin (bovine) 158 kDa/5.29nm; ovalbumin (chicken) 44 kDa/3.05nm; myoglobin (horse) 17 kDa/2.04nm; and vitamin B_12_ 1.35 kDa/0.85nm.

### Size cut-off filtration

Monomer fraction from SEC was passed through a 100kDa MWCO filter (Corning) as instructed by the manufacturer (centrifuged at 15,000 × g for 15 mins at 4°C). Filtered material was immediately collected and protein concentration was determined. The filtrate was aliquoted and frozen in −80°C.

### FRET flow cytometry

Cells were harvested with 0.05% trypsin and fixed in 2% paraformaldehyde (Electron Microscopy Services) for 10min, then resuspended in flow cytometry buffer. The MACSQuant VYB (Miltenyi) was used to perform FRET flow cytometry. To measure CFP and FRET, cells were excited with a 405nm laser, and fluorescence was captured with 405/50nm and 525/50nm filters, respectively. To measure YFP, cells were excited with a 488nm laser and fluorescence was detected with a 525/50nm filter. To quantify FRET, we used a gating strategy similar to that previously described {Holmes: 2014} Percent FRET positivity was used to determine the fraction of cells containing aggregates. For each experiment, ~10,000 cells were analyzed in triplicate. Analysis was performed using FlowJo v10 software (Treestar).

### Protease digestion

Pronase (Roche) was diluted in PBS to a final concentration of 1 mg/mL and single-use aliquots were stored at −80°C. Clarified cell lysate was prepared as previously described and protein concentrations were normalized to 4 μg/μL, unless otherwise noted. 40 μg (10 μL) of cell lysate was added to 10 μL of pronase at a concentration of 60 μg/mL (diluted in PBS) for a final volume of 20 μL and a final pronase concentration of 30 μg/mL. Cell lysates were digested at 37°C for 90min. Reactions were quenched by addition of 20 μL of 2x sample buffer (final SDS concentration of 4%) and boiling for 5 minutes. 15 μL of each sample was loaded onto a 10% Bis-Tris NuPAGE gel (Novex by Life Technologies) and run at 150 V for 65 minutes. Protein was transferred to Immobilon P (Millipore) using a semi-dry transfer apparatus (Bio-Rad) and membranes were probed for tau RD as described above.

### Sedimentation Analysis

Clarified cell lysate was prepared as described previously. 10% of the lysate was set aside as the total fraction; the rest was centrifuged at 186,000 x g for 1 h. The supernatant was placed aside and the pellet was washed with 1 mL PBS prior to ultracentrifugation at 186,000 x g for 30 minutes. The supernatant/wash from this step was aspirated and the pellet was re-suspended by boiling in RIPA buffer with 4% SDS and 100 mM DTT. Bradford assay (Bio-Rad) with BSA standard curve was used to normalize all protein concentrations. 1 μg of total protein was loaded per well on a 4-12% Bis-Tris gel (Invitrogen). For all samples 1:1 ratio of supernatant to pellet was used. Gels were analyzed by western blot.

### Confocal microscopy

96-well plates (Costar 3603) were coated with 1X Poly D-Lysine (PDL), incubated overnight at 37°C. Plates were then washed with PBS and cells were plated and grown in DMEM media for 24 h. Media was then removed and replaced with 4% PFA for 10 min. PFA was removed and washed 2x with PBS followed by staining with DAPI for 10 min in 0.05% Triton-X. Cells were washed and stored in PBS. The plate was imaged with In Cell Analyzer 6000 at 40x resolution with the assistance of the UTSW High Throughput Screening core facility. Images were coded and a blinded counter scored aggregate morphology, blinded to conditions.

### Cell lysate production for animal inoculation experiments

The cell lines were grown in 10 cm dishes until 80% confluency. The cells were then washed, trypsinized, resuspended in media and centrifuged at 1000 × g. Cell pellets were washed with PBS. The pellet was then stored at −80°C. Prior to analysis, pellets were thawed on ice and re-suspended in 1× PBS with cOmplete protease inhibitors (Roche) and sonicated using an Omni-Ruptor 250 probe sonicator at 40% power for 35 × 3 second cycles. The probe sonicator was washed with 10% bleach, 100% ethanol and ddH_2_O between cell lines. Strains were subsequently centrifuged at 1000 × g, normalized to 7 μg/uL by Bradford assay (Bio-Rad) and stored in aliquots at −80°C.

### Animal maintenance

We obtained transgenic mice that express 4R1N P301S human tau under the murine prion promoter (5) from Jackson Laboratory, and maintained them on a B6C3 background. Transgenic mice and wild-type littermates were housed under a 12 hour light/dark cycle, and were provided food and water *ad libitum*. All experiments involving animals were approved by the University of Texas Southwestern Medical Center institutional animal care and use committee.

### Inoculation of mouse brain

P301S mice were anesthetized with isoflurane and kept at 37°C throughout the inoculation. Mice were injected with separate 10 μL gas-tight Hamilton syringes for each strain at a rate of 0.2 μL per minute. Animals were inoculated with 10 μg (1.428 μL) of cell lysate in the left hippocampus (from bregma: −2.5 mm posterior, −2 mm lateral, −1.8 mm ventral). Animals were then allowed to recover and monitored for 40 days prior to tissue collection.

### Animal tissue collection

P301S or WT mice were anesthetized with isoflurane and perfused with chilled PBS + 0.03% heparin. Brains were post-fixed in 4% paraformaldehyde overnight at 4°C and then placed in 30% sucrose in PBS until further use.

### Histology

Brains were sectioned at 50μm using a freezing microtome. Slices were first blocked for one hour with 5% non-fat dry milk in TBS with 0.25% Triton X-100 (blocking buffer). For DAB staining, brain slices were incubated with biotinylated AT8 antibody (1:500, Thermo Scientific) overnight in blocking buffer at 4°C. Slices were subsequently incubated with the VECTASTAIN Elite ABC Kit (Vector Labs) in TBS prepared according to the manufacturer’s protocol for 30 minutes, followed by DAB development using the DAB Peroxidase Substrate Kit with the optional nickel addition (Vector Labs). Slices were imaged using the Olympus Nanozoomer 2.0-HT (Hamamatsu).

## Acknowledgements

We thank Sushobhna Bhatra, Dana Dodd, Lukasz Joachimiak, Hilda Mirbaha, Victor Manon, William Russ, Barbara Stopschinski, Jaime Vaquer-Alicea, Amy Zwierzchowski-Zarate for critiques. This work was supported by grants from the Tau consortium and NIH grants awarded to 1R01NS071835 (M.I.D), R01NS089932 (M.I.D). We appreciate the help of High Throughput Screening Core (HTS) administered by Bruce Posner, PhD. Human tissue samples were provided by the Neurodegenerative Disease Brain Bank at the University of California, San Francisco.

## References

1. Neurodegenerative tauopathies. 2001;24(1):1121–59. Retrieved from: http://www.annualreviews.org/doi/abs/10.1146/annurev.neuro.24.1.1121

2. Frost B, Ollesch J, Wille H, Diamond MI. Conformational diversity of wild-type Tau fibrils specified by templated conformation change. J. Biol. Chem. American Society for Biochemistry and Molecular Biology; 2009 Feb 6;284(6):3546–51. PMCID: PMC2635036

3. Frost B, Jacks RL, Diamond MI. Propagation of tau misfolding from the outside to the inside of a cell. J. Biol. Chem. American Society for Biochemistry and Molecular Biology; 2009 May 8;284(19):12845–52. PMCID: PMC2676015

4. Clavaguera F, Bolmont T, Crowther RA, Abramowski D, Frank S, Probst A, et al. Transmission and spreading of tauopathy in transgenic mouse brain. Nat. Cell Biol. 2009 Jul;11(7):909–13. PMCID: PMC2726961

5. Yoshiyama Y, Higuchi M, Zhang B, Huang S-M, Iwata N, Saido TC, et al. Synapse loss and microglial activation precede tangles in a P301S tauopathy mouse model. Neuron. 2007 Feb 1;53(3):337–51.

6. Sanders DW, Kaufman SK, Devos SL, Sharma AM, Mirbaha H, Li A, et al. Distinct Tau Prion Strains Propagate in Cells and Mice and Define Different Tauopathies. Neuron. 2014 May 21. PMCID: PMC4171396

7. Clavaguera F, Akatsu H, Fraser G, Crowther RA, Frank S, Hench J, et al. Brain homogenates from human tauopathies induce tau inclusions in mouse brain. Proc. Natl. Acad. Sci. U.S.A. National Acad Sciences; 2013 Jun 4;110(23):9535–40. PMCID: PMC3677441

8. Fitzpatrick AWP, Falcon B, He S, Murzin AG, Murshudov G, Garringer HJ, et al. Cryo-EM structures of tau filaments from Alzheimer’s disease. Nature. 2017 Jul 5;56:343. PMCID: PMC5552202

9. Kaufman SK, Sanders DW, Thomas TL, Ruchinskas AJ, Vaquer-Alicea J, Sharma AM, et al. Tau Prion Strains Dictate Patterns of Cell Pathology, Progression Rate, and Regional Vulnerability In Vivo. Neuron. 2016 Nov 23;92(4):796–812. PMCID: PMC5392364

10. Tau Trimers Are the Minimal Propagation Unit Spontaneously Internalized to Seed Intracellular Aggregation. 2015 Jun 12;290(24):14893–903. PMCID: PMC4463437

11. Ohhashi Y, Yamaguchi Y, Kurahashi H, Kamatari YO, Sugiyama S, Uluca B, et al. Molecular basis for diversification of yeast prion strain conformation. Proc. Natl. Acad. Sci. U.S.A. National Academy of Sciences; 2018 Mar 6;115(10):2389–94.

